# Clarification and confocal imaging of the non-human primate placental micro-anatomy

**DOI:** 10.1101/330647

**Authors:** James A. Sargent, Victoria HJ Roberts, Jessica Gaffney, Antonio E. Frias

**Author notes:** Corresponding author (JS). James Sargent, Victoria Roberts, and Antonio Frias contributed equally to this manuscript. Jessica Gaffney made a significant contribution to this manuscript.

## Abstract

Placental function is essential for the development of the fetus, and is – in part – related to the 3D arrangement of the villous and vascular geometry. Recent advances in tissue clarification techniques allow for deep high-resolution imaging with confocal microscopy without altering the spatial characteristics of the tissue. These image stacks can be analyzed quantitatively to provide insights regarding the villous and vascular micro-anatomy as well as the interrelationships between the two. However, such analyses require optimization of the tissue preparation, immuno-labeling, and clarification protocol in order to provide reliable results suitable for the detection of subtle differences in pathologic pregnancies. Placental and fetal development are similar between human and non-human primate pregnancies, with the latter serving as a reliable, validated, highly-controlled, well-characterized translational model for the former.

We present a protocol for the preparation, immuno-labeling, and clarification of the non-human primate placenta optimized for confocal microscopy and subsequent quantification of the micro-anatomic structures.

## 1. Introduction

To ensure adequate gas exchange and transportation of nutrients to the fetus, the appropriate development of the maternal and fetal placental vascular networks is essential [1-2]. The caliber, branching patterns, and interrelationships of the fetal vasculature and the terminal and stem villi are thought to be essential for maintaining adequate uniform blood supply throughout the placenta. Dysfunctional alterations in these morphologies or relationships have been linked to post-placental hypoxia and subsequent intrauterine growth restriction, hypertensive disorders of pregnancy, and stillbirth [1-8].

Until recently, assessment of the placental vasculature has largely relied upon stereological or vascular casting methods. Traditional stereology utilizes paraffin-embedded tissue slices and reproducible counting methods to estimate variables such as villous or vessel size, number, surface area and volume; more complex computational methods can then be employed to estimate diffusion distance and complex hemodynamic parameters [8-11]. While stereology allows the observer to assess for additional histologic variables (e.g. syncytial knots, placental infarcts, and villous mal-development) it is limited to a 2D view without preserving the 3D vascular and villous geometry [1,2,12]. To address these issues, corrosion casting of the placental vasculature was developed as a means to preserve the vascular morphology for qualitative and quantitative assessment [13-17]. Unfortunately, these methods are technically complex, they can lead to distortion or destruction of the fetal capillaries, and because of the corrosive tissue digestion process, they eliminate the potential to evaluate the relationship between the fetal vascular bed and the surrounding villous tissue [14]. As such, new methods are required that allow for the investigation of the 3D vascular structure and the villous architecture.

Through the imaging of immuno-fluorescently labeled and clarified tissue with a confocal microscope, new approaches are emerging that address the shortfalls of stereological and vascular casting methods. The use of confocal microscopy to capture precisely-located optical sections allows for the 3D rendering of the images [18-20], however the depth of tissue analysis has been limited due to refractive index mismatching and subsequent light-scattering [21]. The clarification of biologic specimens through refractory index equilibration has been studied since the beginning of the 20^th^Century, however earlier methods were damaging to tissue and incompatible with fluorescently-labeled specimens [22]. To take advantage of the increasingly sophisticated confocal methodologies available, newer tissue clarification methods have been developed that are either based on hydrogel embedding, hyper-hydration, simple immersion, or solvent-based techniques; each of these having their respective deleterious effects on tissue structure or compatibility issues with immunofluorescently labeled tissue [22]. Visikol^®^HISTO™ is a proprietary reversible clearing technique that uses an ethanol-based solvent, which causes minimal tissue alteration (shrinkage/expansion of <5%) and is compatible with immuno-fluorescent staining. Importantly, the tissue is preserved in a way that can be reversed, thus allowing for post-confocal fixation and paraffin embedding; this facilitates validation with traditional stereological methods. Recently, Visikol^®^HISTO™ has been utilized to generate 3D renderings of immunofluorescently labeled and clarified human placental tissue [23,24], and this has generated interest in customizing its use for novel analyses.

Numerous computer programs have been created to process confocal images and allow for image optimization, 3D rendering, and subsequent analysis. ImageJ2 is a versatile, java-based, image-processing, free software developed by the NIH which has a flexible development model allowing for simple user-generated plug-ins to increase the programs functionality [18]. These validated, reliable functionalities, and the abundance of online and print tutoring resources, have led to ImageJ2 being used extensively across a diverse range of scientific fields [25,26]. The 3D image-based quantification of vascular networks has been approached in various fields of medicine [27], and ImageJ2 has a multitude of plugins which have been validated for this use [28].

Placental and fetal development are similar between humans and non-human primates (NHP), and NHPs have been used successfully as a translational model to human pregnancies [29]. Numerous, validated, well-characterized, highly-controlled NHP pregnancy disease models exist providing a unique opportunity to assess underlying disease mechanisms and potential therapeutic strategies. Using these methods, quantified analysis of the NHP placenta has not yet been accomplished, however this could provide critical insights in both human and NHP pregnancies. Our objective was to develop and optimize a protocol for the immunofluorescent labeling and clarification of the placenta in the NHP to allow for high-resolution 3D imaging of sufficient quality for advanced quantification methods (Figure 1).

**Figure 1:**
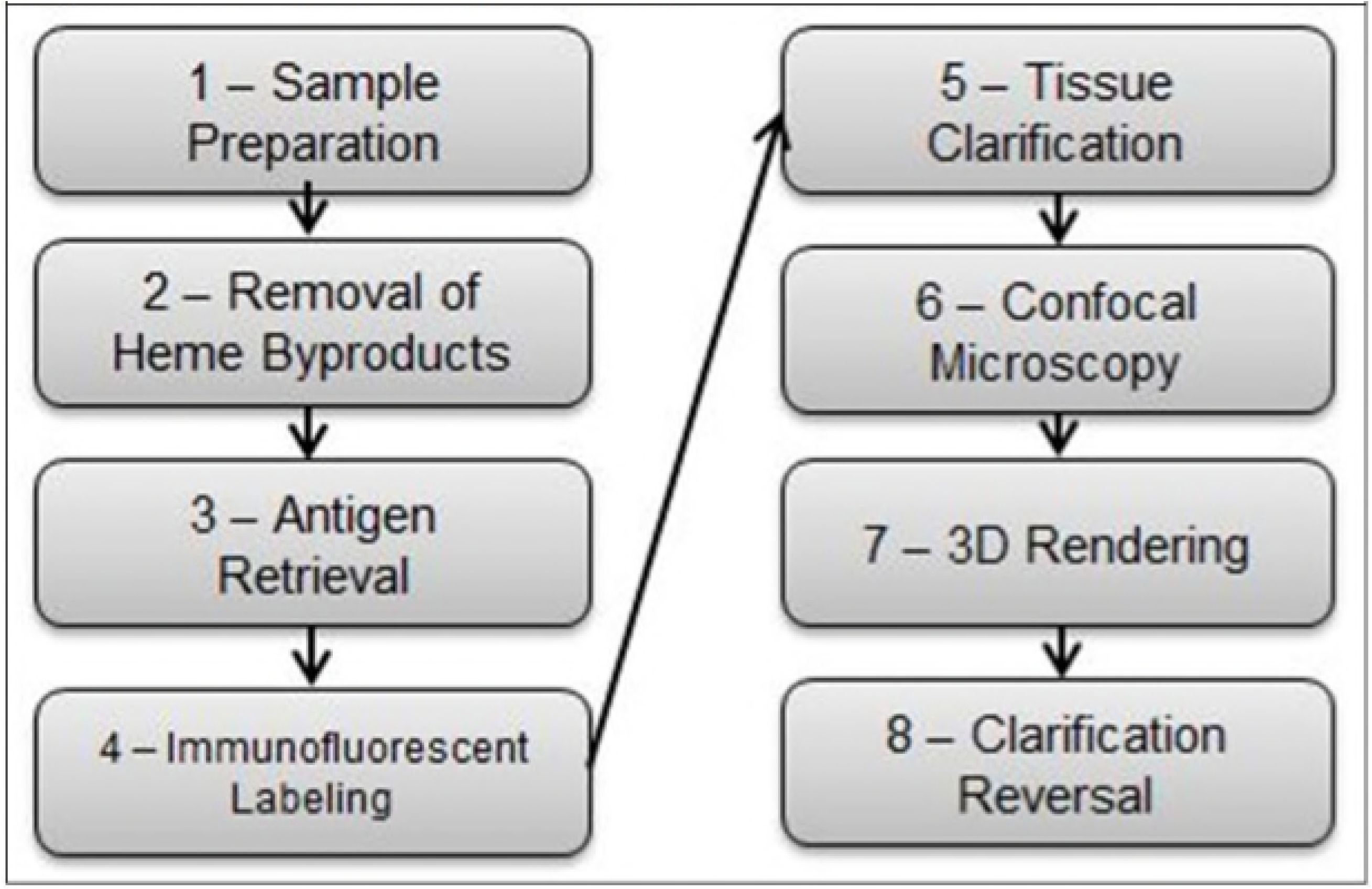
Workflow for Protocol

## 2. Materials and Methods

### 2.1 Sample Preparation

All animal procedures were conducted in accordance with the guidelines of the Institutional Animal Care and Use Committee (IACUC) of the Oregon National Primate Research Center (ONPRC). The ONPRC abides by the Animal Welfare Act and Regulations enforced by the United States Department of Agriculture. Adult female rhesus macaques underwent time-mated breeding to generate pregnancies from which placental tissue was obtained. All animals were pair housed and maintained on a control standard chow diet with additional access to daily enrichment foods and foraging devices. All placental tissues were obtained at the time of cesarean section delivery at gestational day 140 (term being 168 days), immediately placed in PBS, and the decidua and membranes dissected away from the villous tissue. 1mm^3^tissue sections were isolated and immersed in 10% Zinc Formalin for 4 hours prior to transfer in 70% ethanol and storage at 4°C.

### 2.2 Removal of Heme Byproducts

To address background fluorescence caused by hemoglobin, fixed tissue was placed in PBS and gently rocked at 4°C (Boekel Rocker II 260350-2 Platform Rocker, Boekel Scientific Feasterville, PA) for 24 hours. Tissue was dehydrated with 50%, 75%, and 100% methanol washes with gentle rocking at 4°C for 10 minutes per wash. Using a bleaching solution (30% H_2_O_2_, DMSO, methanol in a 1:1:4 ratio), tissue was incubated at 4°C for 24 hours under gentle rocking. To reduce auto-fluorescence, heat exposure was limited and all steps were performed under LED bulbs with wavelengths of 480nm and 660nm (45W Multi-Spectrum LED, iPower, Irwindale, CA) at a distance of 6cm.

### 2.3 Antigen Retrieval

For immunofluorescent staining, antigen retrieval was essential. Tissue was rehydrated using 20% DMSO/methanol, 75% and 50% methanol, and PBS washes with gentle rocking at 4°C for 10 minutes per wash. Optimal antigen retrieval was obtained using Citrate Buffer (10mM Citric Acid with 0.05% Tween at pH 6) exposed to 80°C at 6 psi (Cuisenart Electric Pressure Cooker – CPC600, Cuisenart, East Windsor, NJ) for 40 minutes, and then placed at 21°C for 20 minutes.

### 2.4 Immunofluorescent Labeling

Blocking and permeabilization was performed using a solution of 2% Donkey Serum and 0.2% Triton X-100 in PBS with gentle rocking at 4°C for 24 hours under LED exposure at 4°C. A primary antibody solution of a monoclonal mouse-anti-human CD31 (Thermofisher MA1-26196, endothelial cell marker) and a polyclonal rabbit-anti-human CK7 (Abcam, ab103687, trophoblast marker) was made at 1:10 concentrations in a solution of 2% Donkey Serum in PBS, and tissue was immersed and placed at 37°C for 72 hours. Four 10 minute washes of 2% Donkey Serum in PBS at 21°C were then performed before placing in a secondary antibody solution of donkey-anti-mouse pre-adsorbed polyclonal antibody (Abcam, ab150109, alexafluor 488) and a donkey-anti-rabbit antibody (Abcam, ab150075, alexafluor 647) at a concentration of 1:250 for both secondaries in 2% donkey serum and PBS for 48 hours at 21°C with agitation at 150rpm.

### 2.5 Tissue Clarification

Samples were washed four times in 2% donkey serum and PBS for 10 minutes at 21°C at 150rpm prior to dehydration. Three 10-minute washes of 50%, 75%, and 100% methanol were performed before immersing the tissue in the Visikol^®^Histo1^™^for 4 hours at 21°C at 150rpm. The tissue was placed in the Visikol^®^ Histo2^™^ solution for a minimum of 4 hours, and storage at 21°C.

### 2.6 Confocal Microscopy

Clarified samples were placed in a Sykes-Moore chamber (Bellco Glass Inc, Product 1943-11111, Vineland, NJ), between two 25mm glass coverslips, and immersed in 650 uL of Visikol^®^HISTO2™ solution. Imaging was performed using a Leica SP5 AOB5 spectral confocal system in two-channel (Leica/ALEXA 488, Leica/ALEXA 633) optical sections at optimized intervals calculated by the imaging system (∼1µm per slice, 200-300 slices per sample) using a x20 PL APO NA 0.75 IMM CORR CS2 air objective. Optical detectors were set to 430-480nm and 635-680nm respectively to minimize crossover. To increase resolution, the confocal was set to 1024×1024 pixels, pinhole of 0.4 Airy, at 200 Hz acquisition speed, with line averaging. Negative controls (no secondary antibody) were used to adjust the confocal gain sensitivity to eliminate auto-fluorescence.

### 2.7 3D Rendering

Imaged z-stacks were uploaded into Fiji-ImageJ2 as hyperstacks and the stacks were projected into either maximum intensity or 3D renderings.

### 2.8 Clarification Reversal and Validation with Microscopy

Post-imaging, tissue samples were immersed in 100% ethanol at 21°C until opacified, and then stored in 70% ethanol. Tissues were dehydrated with increasing concentrations of ethanol before being mounted in paraffin, sectioned, and stained for H&E, CD31, and CK7. Slides were imaged using a Leica Leitz Diaplan microscope (Leitz Wetzlar, Germany) using the PL Fluotar 16x / NA 0.45 and NPL-Fluotar 40x / NA 0.70 objectives.

## 3. Results and Discussion

We describe the developed protocol for the successful immuno-fluorescent labeling and clarification of the placenta of the NHP, which has been optimized for confocal microscopy and quantitative analysis. We approached protocol development as an iterative process with the aim to maximize both reproducibility and the signal-to-noise ratio of our immuno-fluorescent labeling as well as to minimize the need for post-imaging processing. Our pre-determined threshold for resolution was the diameter of the capillary vessels – 3µm – and z-stacks collected at 20x magnification allow for clear delineation of the capillary vessels within the terminal villi (Figure 2 & 3).

**Figure 2:**
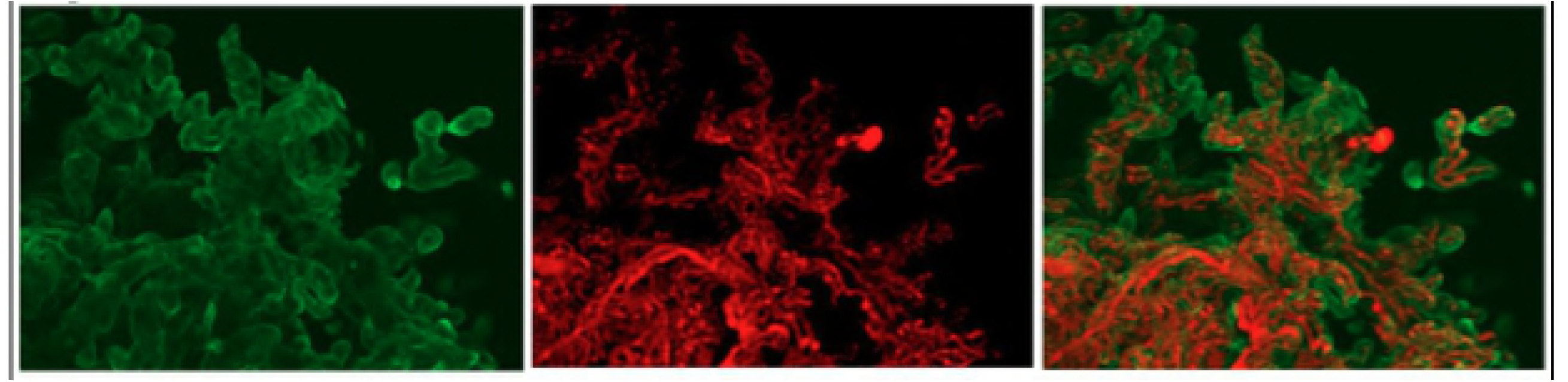
Maximum-intensity projections of immuno-fluorescent labeled and clarified NHP placental tissue imaged using the Leica SP5 AOB5 spectral confocal system with a 20x PL APO NA 0.75 AIR CORR CS2 objective at 647nm (left), 488nm (center), and then overlayed (right).

**Figure 3:**
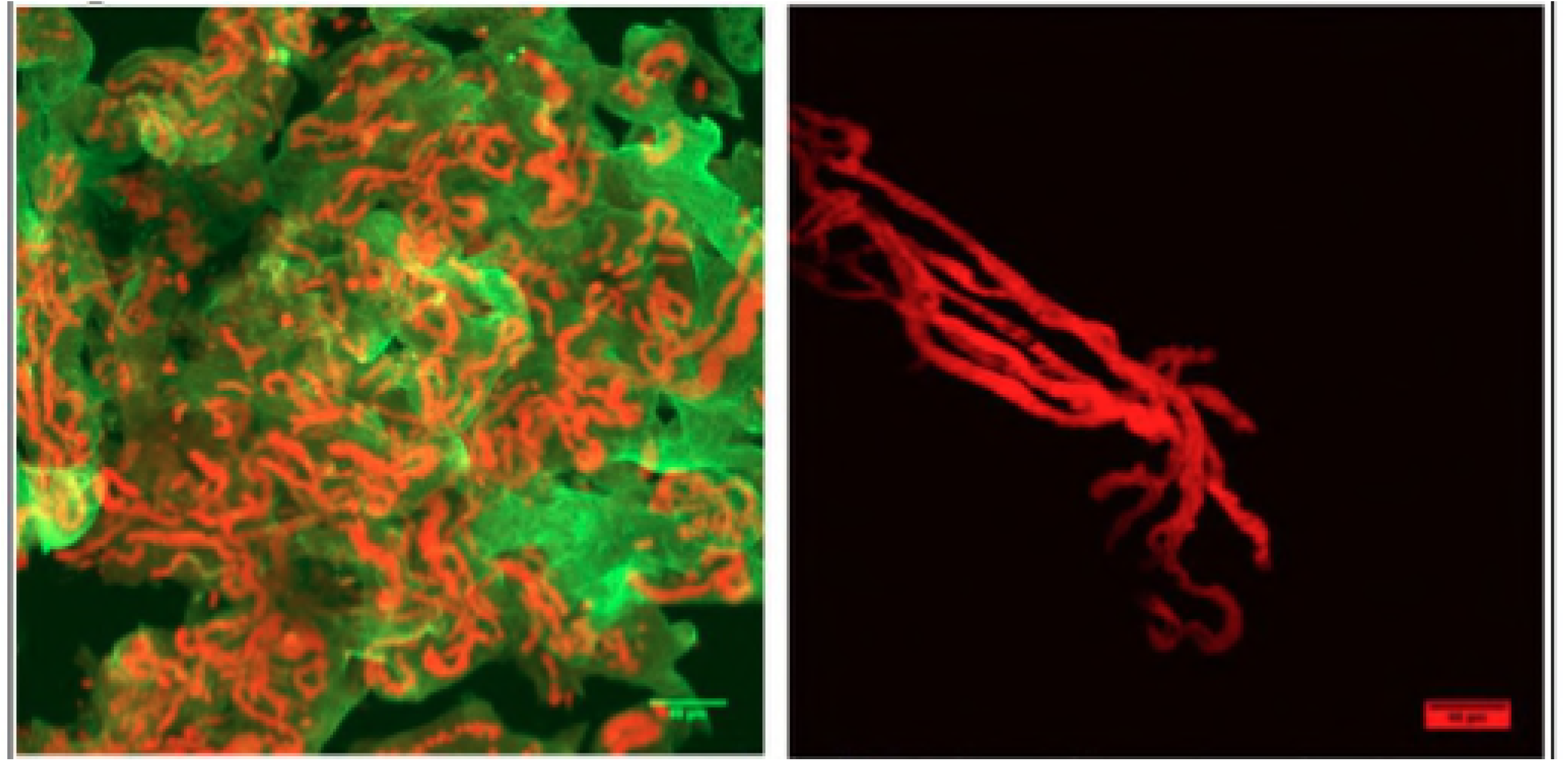
Maximum-intensity projections of clarified NHP placental tissue imaged at 40x magnification with immuno-labeling of CD31 (red) and CK7 (green) antigens.

### 3.1 Experimental Considerations

#### 3.1.1 Auto-fluorescence

Fixed placental tissue, in the absence of immune-labeling, has an abundance of natural auto-fluorescent noise making it unsuitable for quantitative analysis (Figure 4). Interfering auto-fluorescence can come from multiple sources including naturally occurring organic molecules, retained heme byproducts, and they can also be induced by the fixation process. While imaging techniques and post-imaging correction methods exist, these are unreliable and can reduce the desired signal from immuno-labeled structures. For this protocol, duration of Zn Formalin exposure was minimized to decrease the amount of fixation-induced auto-fluorescence. Clearance of red blood cells can be achieved through saline perfusion of fresh placenta, however this is technically difficult in the NHP and not always feasible. Rinsing the tissue with PBS followed by a H_2_O_2_-based bleaching solution under optimized conditions reliably removed speckling from retained heme byproducts. Lastly, minimizing heat exposure and exposing the tissue to LED Photobleaching further decreased the background natural and fixation-induced auto-fluorescence.

**Figure 4:**
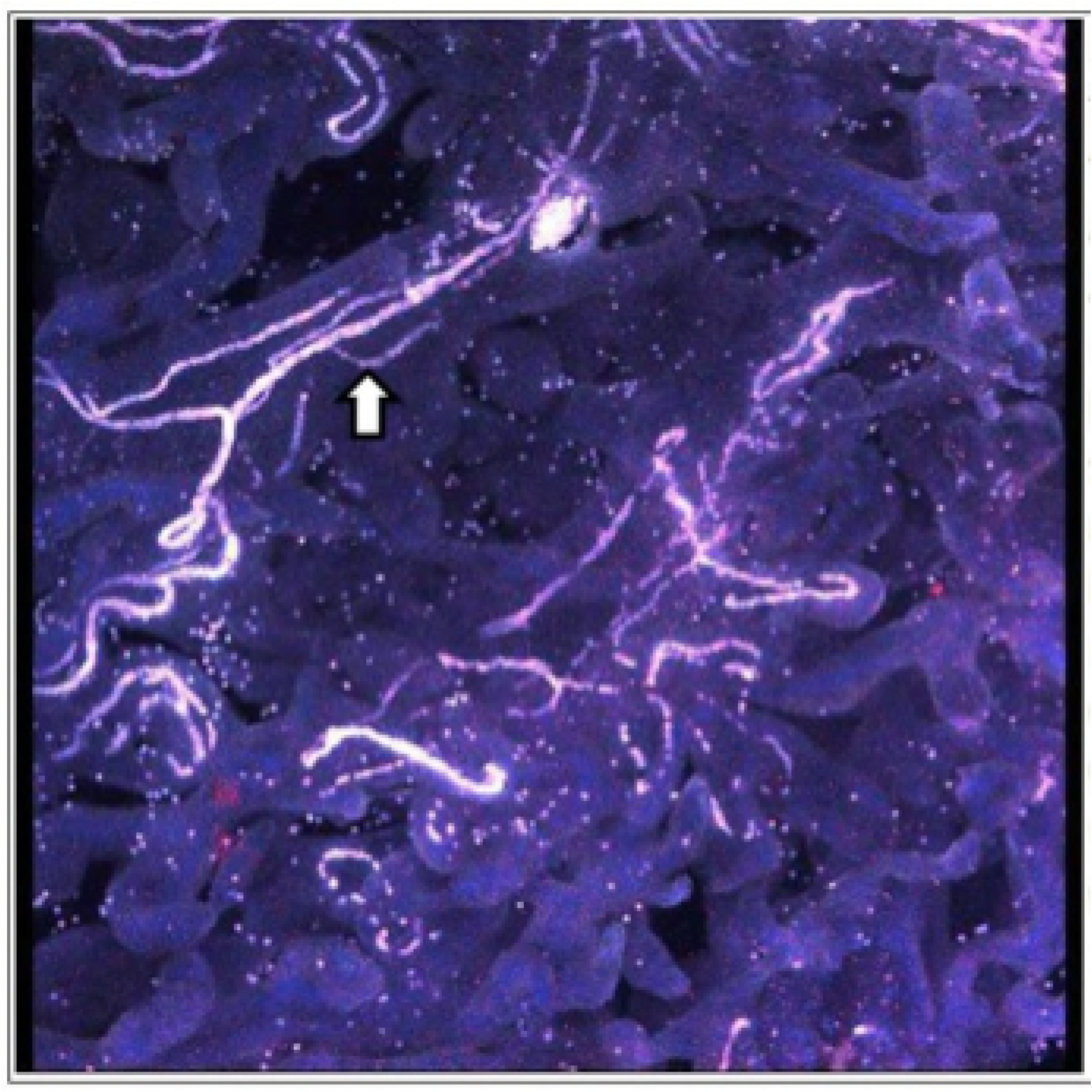
Maximum-intensity projection of fixed, unlabeled, non-clarified placental tissue of the NHP imaged using the Leica SP5 AOB5 spectral confocal system with a 20x PL APO NA 0.75 AIR CORR CS2 objective at 512nm. Arterioles (white arrow) are high-intensity due to elastin auto-fluorescence while hemoglobin byproducts are visualized as “Christmas lights” throughout the image. Villous tissue is visible secondary to natural auto-fluorescence from lipofuscin and other organic molecules. The gradual fading of the intensity, most apparent in the upper left corner of the image, reflects the depth-dependent light-scattering.

#### 3.1.2 Antigen Retrieval

We found antigen retrieval to be necessary for our CD31 and CK7 antigens, with the temperature, duration, and pH optimized as a balance between the two antibodies. In addition, numerous alternative primary antibodies and lectins were trialed under varying conditions (concentration, agitation, duration, temperature) and antigen retrieval steps (buffer type, pH, duration, temperature, pressure settings). Prolonged, high-concentration washes under heat were required to obtain optimal signal strength and tissue penetration.

#### 3.1.3 Clarification and Reversal

Tissue clarification was carried out as per the manufacturer recommendations. While the use of the second solution (Visikol^®^Histo2^™^) was listed as being optional for vascular tissues like the placenta, omitting this step dramatically decreased the effectiveness of the clarification process. Reversal of the clarification process and subsequent H&E stained paraffin-embedded slides revealed no gross morphologic alterations when compared with formalin-fixed tissue from the same placenta that had not undergone the clarification protocol.

### 3.2 Image Acquisition and Analysis

ImageJ is a freeware program capable of rendering uploaded z-stacks into 3D and subsequently analyzing the geometry of the placental micro-anatomy (Figure 5). While filters exist for the smoothing of immune-labeled structures and subtracting out background noise, we sought to minimize the amount of post-imaging processing required prior to analysis by optimizing the signal-to-noise ratio of our immunofluorescent-labeling and clarification protocol. Prior to image analysis z-stacks must be made binary, a process whereby each voxel is analyzed and those that have an intensity below a specified threshold are removed while those that are above threshold are altered to a uniform intensity. This crucial step requires accurate determination of the set threshold, and z-stacks containing heme byproducts, unevenly stained structures, or increased auto-fluorescence result in the loss of labeled structures and the incorporation of unwanted noise interpreted as structures of interest. Through the optimization of our protocol, ImageJ becomes more sensitive and specific to our immuno-labeled objects of interest and the subsequent quantitative analysis is more robust.

**Figure 5:**
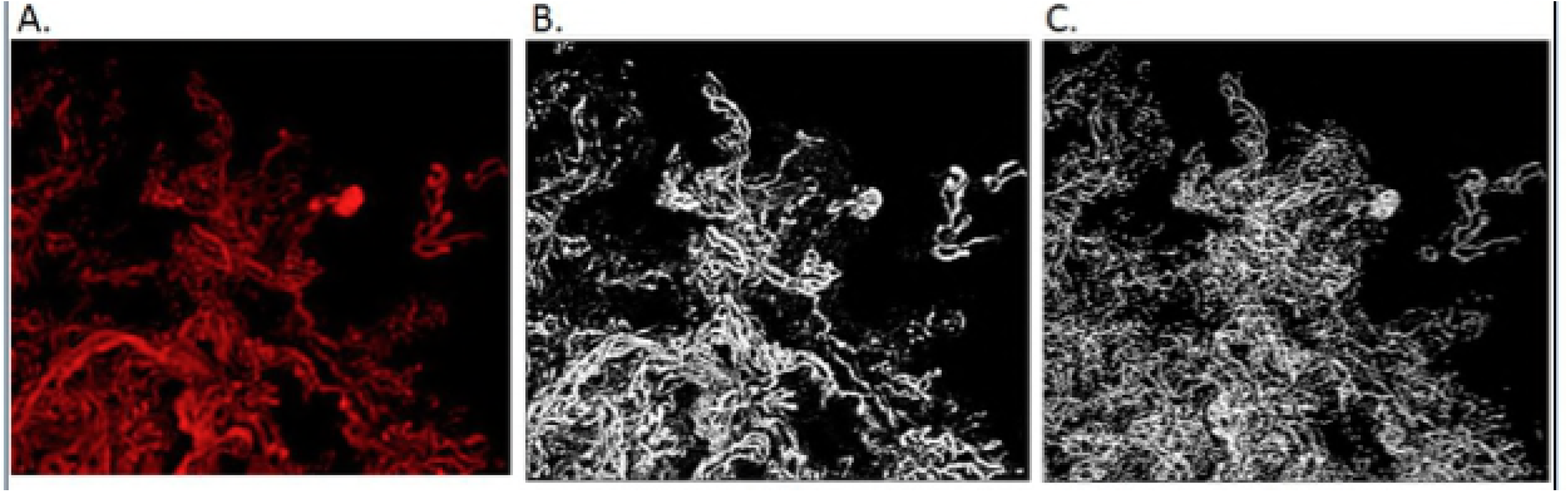
Maximum-intensity projections of CD31 labeled and clarified NHP placental tissue before (A) and after (B) the ImageJ2 thresholding process and skeletonization (C) steps required prior to quantified analysis.

### 3.3 Future Directions

With the development of new analytic tools our ability to determine disease etiology and monitor response to therapy improves. The protocol as described in this paper allows for the generation of confocal imaged z-stacks with sufficient resolution to pursue quantitative analysis.

Modification of this protocol could be used for other 3D analyses including the localization of viral and bacterial antigens, cells of the immune system, membrane proteins and transporters, or genetic alterations (e.g. placental mosaicism).

## Acknowledgements

We would like to thank Eliot Spindel from the Oregon National Primate Research Center, George Merz from the New York State Institute for Basic Research in Developmental Disabilities and Tom Villani from Visikol^®^for their support in developing this protocol.

